# Contribution of linear and nonlinear mechanisms to predictive motion estimation

**DOI:** 10.1101/2021.11.09.467979

**Authors:** Belle Liu, Arthur Hong, Fred Rieke, Michael B. Manookin

## Abstract

Successful behavior relies on the ability to use information obtained from past experience to predict what is likely to occur in the future. A salient example of predictive encoding comes from the vertebrate retina, where neural circuits encode information that can be used to estimate the trajectory of a moving object. Predictive computations should be a general property of sensory systems, but the features needed to identify these computations across neural systems are not well understood. Here, we identify several properties of predictive computations in the primate retina that likely generalize across sensory systems. These features include calculating the derivative of incoming signals, sparse signal integration, and delayed response suppression. These findings provide a deeper understanding of how the brain carries out predictive computations and identify features that can be used to recognize these computations throughout the brain.

## INTRODUCTION

Sensory regions of the brain provide a window to the outside world, allowing animals to infer information about the external environment and, ultimately, to interact with that environment. A central tenet of sensory neuroscience, the notion of feature selectivity, states that neuronal responses depend on a relatively small number of features present in the incoming stimulus (Fairhall et al., 2006; Sharpee et al., 2004; Pillow and Simoncelli, 2006; Barlow et al., 1964; Zhang et al., 2012; Hubel and Wiesel, 1959). Indeed, there is strong evidence that the brain has evolved the ability to efficiently encode incoming sensory inputs by matching neural response properties to the structure of the natural environment and specifically those aspects of nature with the greatest behavioral relevance (Barlow, 1961; Rieke et al., 1995; Olshausen and Field, 1996; Lewicki, 2002; Machens et al., 2005; Laughlin, 1981; Fairhall et al., 2001; Reinagel, 2001; Machens et al., 2001; Vinje and Gallant, 2002; Chacron et al., 2003; Escabí et al., 2003).

A strong version of this hypothesis further posits that the information most useful for guiding behavior is that information from the past that can be used to estimate future states of the environment—the predictive information (Bialek et al., 2001; Salisbury and Palmer, 2016; Tishby et al., 1999). Predictive encoding in sensory systems is currently best understood in the context of visual motion estimation, where retinal neurons use the past positions of a moving object to estimate its future trajectory (Berry et al., 1999; Johnston and Lagnado, 2015; Leonardo and Meister, 2013; Palmer et al., 2015; Schwartz et al., 2007; Liu et al., 2021). Predictive computations should also be present in other sensory systems (Sachdeva et al., 2021; Bialek et al., 2001; Salisbury and Palmer, 2016; Singer et al., 2018; Chalk et al., 2018). For example, an animal foraging for food must utilize the spatiotemporal patterns of odours in the environment to estimate the location of a food source (Vergassola et al., 2007; Vickers, 2000; Koehl et al., 2001; Zelano et al., 2011). A deeper mechanistic understanding of predictive computations is needed to identify these computations across neural systems.

Here, we combine neural recordings, dimensionality reduction techniques, and neural circuit modeling to identify neural signatures of predictive encoding. We demonstrate that four cell types in the primate retina that show efficient predictive encoding share a common set of low-dimensional features that govern their light responses. These features include both linear and non-linear properties. Several of these features, including the calculation of temporal derivatives, sparse signal integration, and delayed suppression of the neural response, are signatures of predictive encoding that may generalize across sensory systems.

## RESULTS

### Common features of retinal receptive fields

We studied how both linear and nonlinear properties of the spatiotemporal receptive field contribute to motion encoding in On- and Off-type parasol and smooth monostratified ganglion cells in the macaque monkey retina. We focused on these cells because they provide input to brain regions that contribute to motion processing in primates and they efficiently encode predictive motion information (Rodieck and Watanabe, 1993; Crook et al., 2008; Schiller et al., 1990; Billington et al., 2011; Liu et al., 2021). To estimate their receptive-field properties, we recorded spike responses in these cells to a spatiotemporal noise stimulus consisting of adjacent bars presented over the receptive field center and surround regions (grid size, 19 × 1; bar width, 50 μm; height, 730 μm). The contrast of each bar was drawn from a Bernoulli distribution on each stimulus frame (contrast, ±50%; see Methods).

The stimulus set used included stimuli with spatiotemporal correlations and stimuli lacking net correlations (Liu et al., 2021). However, the nature of the spatiotemporal correlations precluded the use of classical dimensionality reduction techniques. Maximally informative dimensions, an information-theoretic technique, does not suffer from this limitation and we utilized this method to estimate the spatiotemporal filtering properties of each cell (Sharpee et al., 2004; Williamson et al., 2015; Paninski, 2003). This technique calculates the set of spatiotemporal filters or kernels that best preserve information about the stimulus in a cell’s spike outputs (Sharpee et al., 2004; Williamson et al., 2015; Pillow and Simoncelli, 2006; Paninski, 2003). The idea is that a single neuron is insensitive to most of the possible stimuli that can be generated; instead the neuron’s limited stimulus selectivity can be described with a relatively small number of spatiotemporal kernels (**Figure 1**, **Figure S1**). These kernels form a simplified (low-dimensional) description of the spatiotemporal patterns that produce spiking in a cell and thus provide useful insights into the cell’s encoding properties.

**Figure 1.**
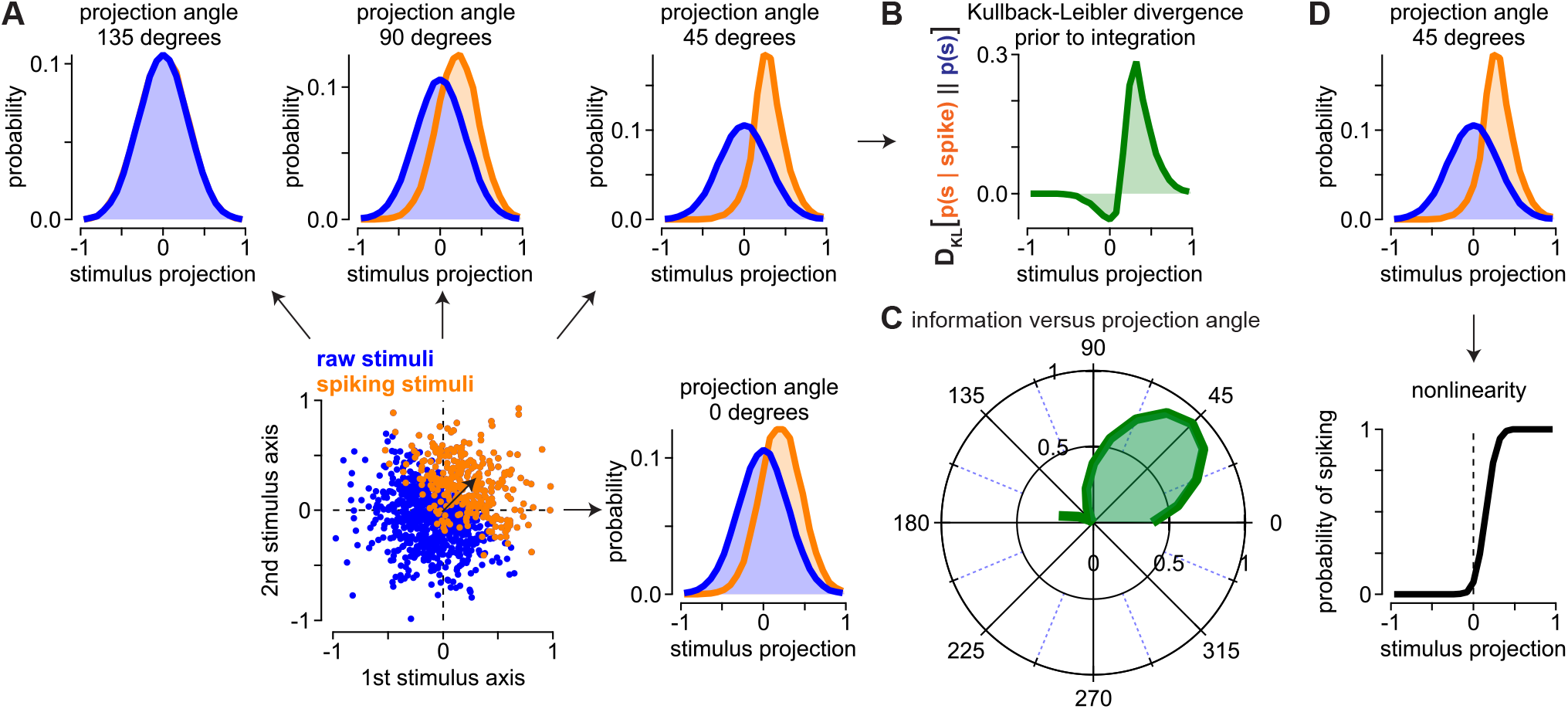
Example of the maximally informative dimensions technique for dimensionality reduction. (A) To illustrate the technique, we generated a hypothetical two-dimensional stimulus space with the blue dots indicating all of the raw stimuli and the orange dots indicating the subset of raw stimuli that elicited a spike. The probability distributions for the raw and spike-triggered stimuli are shown for four projection angles. The distributions showed the greatest separation at 45 degrees, which corresponds to the angle by which the data were artificially rotated. (B) The maximally informative dimensions technique seeks projections that maximize the information that a single spike conveys about the stimulus. This single-spike information is equivalent to the Kullback-Leibler divergence between the spike-triggered and raw stimulus distributions (*green*). Divergence values are shown for the 45 degree projection prior to integration. Integration produces a single value in bits spike^−1^. (C) Polar plot showing the single-spike information as a function of projection angle for the stimulus space in (A). Information peaked at 45 degrees. (D) The probability of observing a spike given a stimulus is equal to the ratio between the spike-triggered and raw stimulus distributions multiplied by the probability of observing a spike.

Our goal was to obtain a low-dimensional representation describing the relationship between the input stimulus and the spike output of each cell. However, the computational overhead of the maximally informative dimensions algorithm is very high, and we were limited to three spatiotemporal kernels in our receptive field estimation (Sharpee et al., 2004; Williamson et al., 2015). For each cell, we computed the three spatiotemporal kernels that preserved the greatest amount of information about the stimulus in the spike output of the cell. The kernels were ordered by their informativeness with the first/dominant kernel preserving the greatest information about the stimulus. These kernels showed consistent spatial features across cells (**Figure 2**). The dominant spatial kernels for all cell types were well approximated by a Gaussian function. The dominant temporal kernels were biphasic and peaked at a time lag of approximately 40 ms, consistent with previous measurements from parasol and smooth monostratified ganglion cells (Rhoades et al., 2019; Pillow and Simoncelli, 2006; Chichilnisky and Kalmar, 2002).

**Figure 2.**
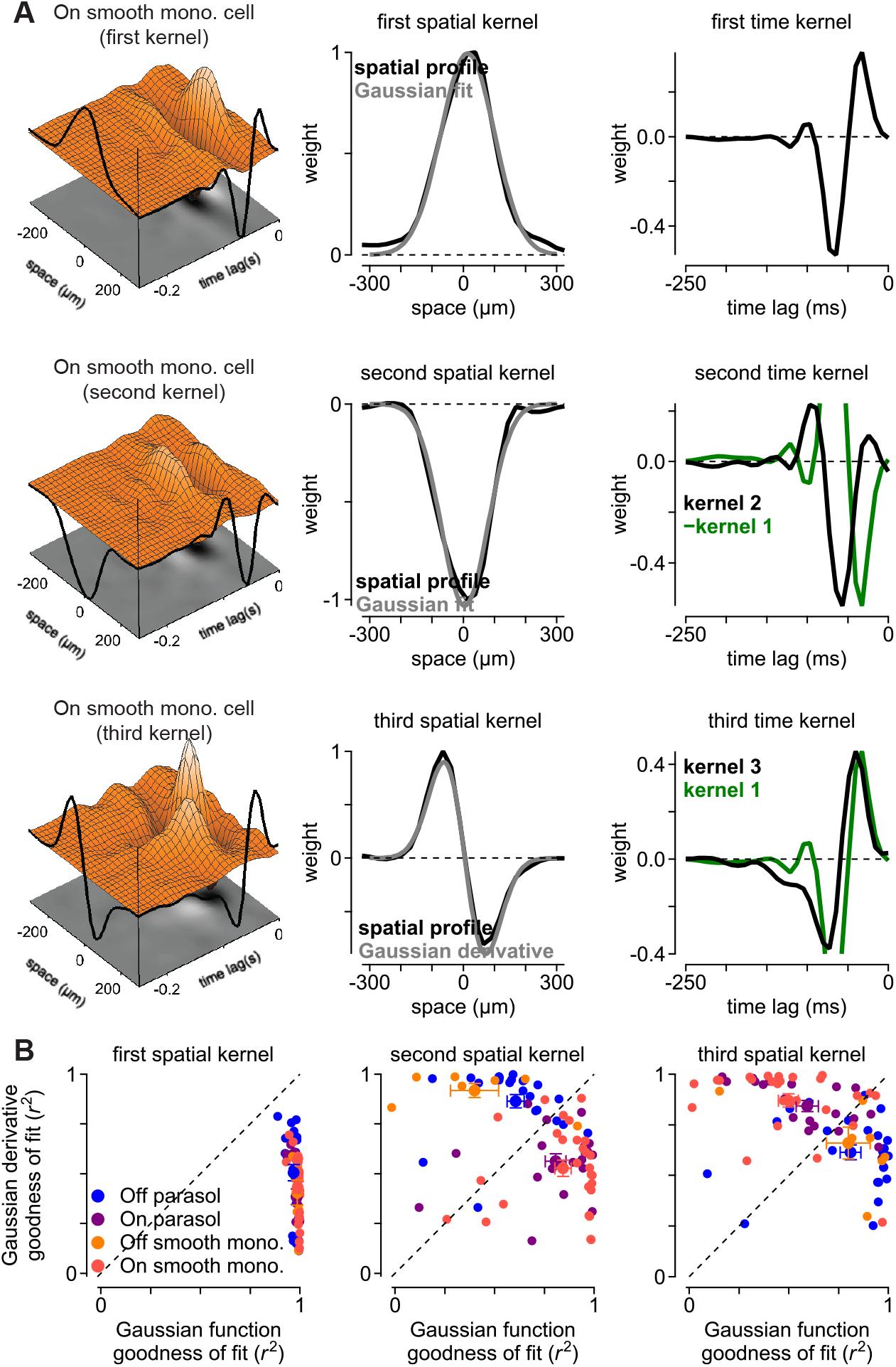
Primate ganglion cells show several significant receptive-field kernels. (A) Spatiotemporal kernels for On and Off smooth monostratified and parasol ganglion cells were determined using an information-theoretic analysis technique. The dominant kernel showed a classical Gaussian spatial profile (*top row*). The spatial profiles of the second (*middle row*) and third kernels (*bottom row*) extracted resembled the first and second derivatives of a Gaussian function, respectively. The scaled temporal component of the first kernel is shown with the second and third kernels to illustrate the differences in kinetics (*green*); the sign of the first kernel was inverted in the second row to match the sign of the second kernel. (B) Goodness-of-fit comparison (*r*^2^) for a Gaussian function versus the first derivative of a Gaussian. The comparison is shown for the first three spatial kernels. Circles and error bars indicate mean ± SEM.

One of the additional kernels showed a spatial profile consistent with the first derivative of a Gaussian function with a positive-going lobe at negative *x*-values and a negative-going lobe at positive *x*-values. This derivative kernel typically occurred as the second kernel in Off-type cells and the third kernel in On-type cells. The other kernel typically occurred as the third kernel in Off-type cells and the second kernel in On-type cells (**Figure 2**B). The temporal kinetics of this kernel were delayed relative to the other two kernels. This delay was approximately 20 milliseconds relative to the first kernel (time-to-peak re to first kernel, −22.4 ± 3.0 ms; n = 78 cells; p = 9.8 × 10^−10^, Wilcoxon signed rank test).

### Receptive field kernels share a common nonlinearity

The dimensionality reduction technique that we used to estimate the spatiotemporal kernels assumes that outputs of these kernels are summed prior to passing through a common nonlinearity (Sharpee et al., 2004; Williamson et al., 2015). We tested this by comparing the shapes of this shared nonlinearity with the non-linearities computed for a model in which the output of each kernel passed through a separate nonlinearity prior to summation (**Figure 3**; see Methods).

**Figure 3.**
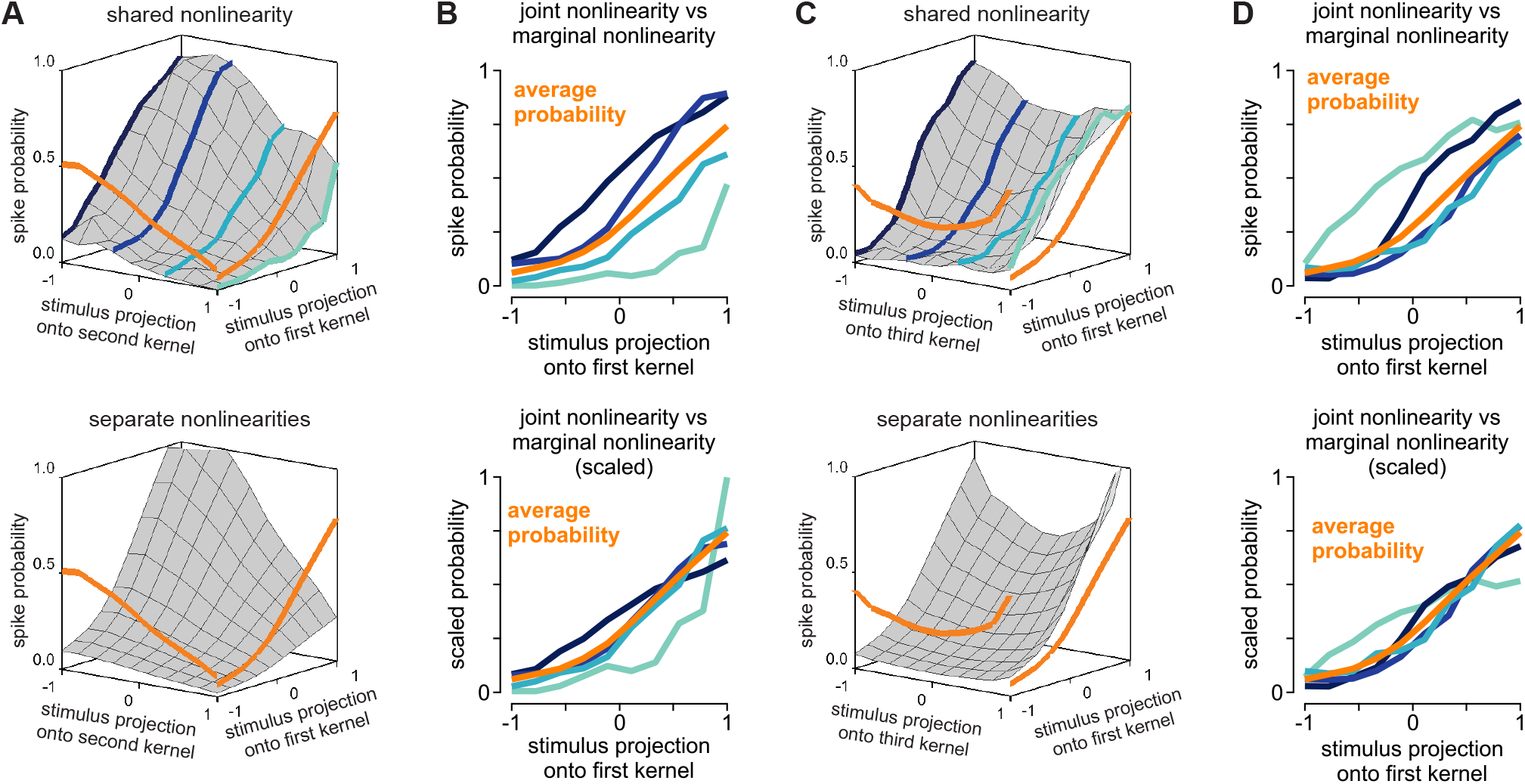
Receptive-field kernels share a common nonlinearity. (A) Two-dimensional nonlinearities illustrating the interactions between the individual kernels for an On smooth monostratified cell. The *x* and *y* axes represent the normalized projection of the stimulus onto the individual kernels. The *z*-axis represents the probability that the cell fired at least one spike. The average spike probabilities for the kernels are shown in orange. The shared nonlinearity was determined by binning the projections and computing the cell’s probability of discharging a spike in each bin (*top*). The separable nonlinearity was calculated from the outer product of the average probability curves and normalizing such that the total spike probability matched that of the shared nonlinearity (*bottom*). (B) Sections through the shared nonlinearity in (A) shown relative to the average probability (*orange*). The sections were multiplied by a scale factor to match to the average probability (*bottom*). The relatively poor match to the average probability indicates that a significant portion of the kernel outputs are combined prior to passing through a shared nonlinearity. (C-D) Two dimensional nonlinearities, as in (A-B), showing interactions between the first and third kernels.

These nonlinearities represent interactions between the kernels in determining the cell’s spike output; both forms of interaction can be captured by computing a two-dimensional surface relating the kernel outputs to the spike response. The *x*-axis and *y*-axis represent the stimulus projections onto the two kernels being examined (**k**_*i*_^⊤^***s***) and the vertical axis shows the spiking probability of the cell for those stimulus projections.

Comparing the two-dimensional nonlinearities illustrates whether the kernels have separate nonlinearities or share a common nonlinearity. If the kernel outputs pass through separate nonlinearities before being combined, then the individual nonlinearities would provide a satisfactory description of spiking behavior in the neuron and the shared and separate nonlinearities would be similar. However, if the kernels shared a common non-linearity, the separately computed nonlinearity would differ from the shared nonlinearity.

Indeed, these nonlinearities differed substantially indicating that the outputs of the kernels were dominated by a single, shared nonlinearity (**Figure 3**). For example, if the kernel outputs passed through separate non-linearities prior to being combined, sections through a particular axis of the two-dimensional nonlinearity should be scaled versions of the average along that axis. However, this was not the case, indicating that a significant proportion of the kernels were combined prior to passing through a common nonlinearity (**Figure 3**B). Thus, these results indicate that the outputs of each of these kernels are combined before passing through a single, dominant nonlinearity (Turner et al., 2018).

### Receptive field modes improve predictive motion encoding

Many dimensionality reduction techniques, including principal components analysis, are technically valid only when the stimulus contrasts are drawn from a Gaussian distribution. Information-theoretic techniques such as the maximally informative dimensions approach employed here do not suffer from this limitation and function properly with non-Gaussian stimuli containing correlations (Sharpee et al., 2004; Pillow and Simoncelli, 2006; Williamson et al., 2015). We confirmed this by recording an uncorrelated stimulus in which the bar contrasts were drawn from a Gaussian distribution; this stimulus was recorded along with our normal stimulus set in the same cell. We then calculated the kernel bases separately for three different stimulus-response sets using the maximally informative dimensions approach: 1) the uncorrelated Gaussian stimulus, 2) the uncorrelated stimulus with bar contrasts drawn from a Bernoulli distribution, and 3) the stimulus set with spatiotemporal correlations included (**Figure 4**). Consistent with theoretical reports, the kernel bases were very similar for the three different stimulus conditions tested—each of the three kernels showed similar spatiotemporal structure across the conditions (Sharpee et al., 2004; Williamson et al., 2015).

**Figure 4.**
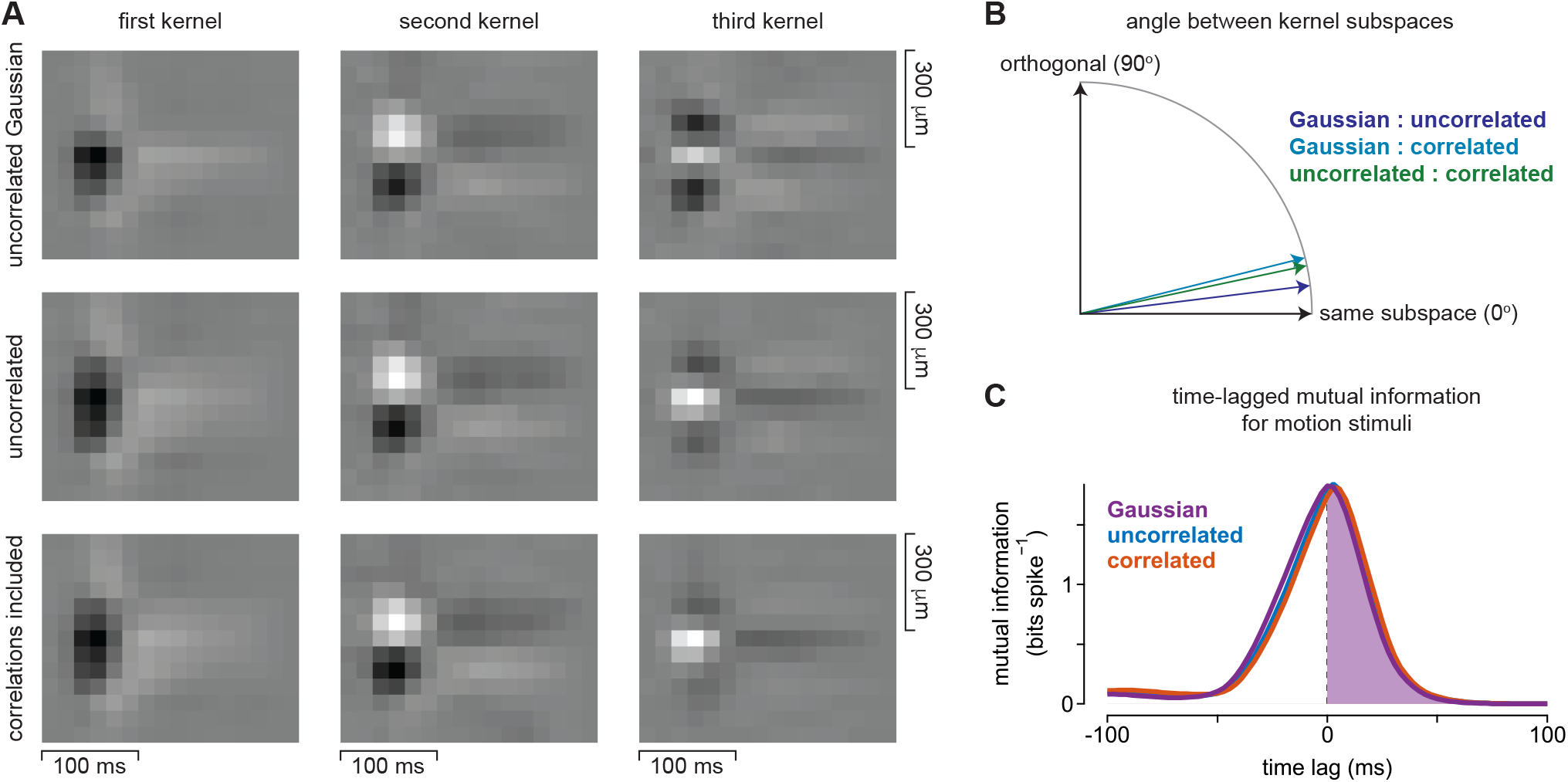
Kernel bases computed with different stimulus classes show similar structures and comparable predictive information encoding. (A) Spatiotemporal kernels in the same cell were estimated for an uncorrelated Gaussian stimulus (*top row*), an uncorrelated stimulus containing contrasts drawn from a Bernoulli distribution (*center row*), and with spatiotemporal correlations included (*bottom row*). The kernel bases computed by the maximally informative dimensions algorithm were similar for the different stimulus classes. (B) Computed angles between the subspaces spanned by the kernel bases in (A). A rotational angle of zero degrees would occur if the kernel bases spanned precisely the same subspace whereas an angle of 90 degrees would occur if the bases were uncorrelated. Rotational angles ranged between 7-14 degrees. (C) Time-lagged mutual information between the stimulus containing pairwise and diverging motion correlations and the output of the kernel bases in (A) computed using the Gaussian stimuli (*purple*), the uncorrelated stimuli (*blue*) or the entire stimulus set which included correlated stimuli (*red*). The shaded regions show the predictive information. The information curves for the kernel bases strongly overlapped, indicating that the subtle differences in bases did not strongly affect predictive motion encoding.

The kernels computed by the maximally informative dimensions algorithm describe a low-dimensional region of stimulus space in which a neuron shows sensitivity to changes in the stimulus features. To determine whether the kernel bases computed for different stimuli defined similar stimulus subspaces, we computed the canonical angles between the kernel bases. An angle of zero degrees occurs when the bases reside on precisely the same subspace and an angle of 90 degrees corresponds to subspaces that are uncorrelated (that is, orthogonal) with each other. The calculated angles between the different kernel bases ranged between 7-14 degrees, indicating that subspaces spanned by the kernel bases were similar but not identical (**Figure 4**B).

To determine whether these differences in the kernels translated to fundamental differences in predictive motion encoding, we computed the time-lagged mutual information for the kernel bases and pairwise and diverging motion correlations that elicited predictive encoding in parasol and smooth monostratified ganglion cells (Liu et al., 2021). This technique measures the information that the spike output of a cell contains about the stimulus at both past and future time lags [(Palmer et al., 2015); see Methods].

The model output was determined by projecting the stimulus onto the kernel basis, summing the kernel outputs, and passing the result through a one-dimensional nonlinearity. This nonlinearity was estimated directly by calculating the spike rate conditioned on the stimulus projection onto the kernel basis (see Methods; **Figure S1**). The resulting mutual information curves strongly overlapped for the computed kernels. The predictive information was also similar for each computed set of kernels (**Figure 4**C, *shaded region*). This result indicates that the slight differences in the estimated kernels do not translate to large differences in predictive motion encoding.

The computed kernels formed a simplified description (i.e., low-dimensional basis) of the spatiotemporal features that best explain the spike responses of these neurons. However, it was not clear whether the additional kernels would improve encoding of predictive motion information relative to the condition in which only the dominant kernel was used. To test this, we projected the motion stimuli onto these kernels and passed the output through the one-dimensional nonlinearity (**Figure 5**). This process was repeated in each cell for four distinct kernel combinations (bases): 1) a basis that comprised only the dominant spatiotemporal kernel, 2) a basis that comprised the first and second kernels, 3) a basis that included the first and third kernels, and 4) a basis that included all three kernels. We then calculated the mutual information between the outputs of these four model bases and the motion stimuli (see Methods). Moreover, we separately calculated the information encoded about the past stimulus (i.e. past information) and future stimulus trajectories (i.e., predictive information). To determine whether the additional kernels improved motion encoding, we normalized the information values relative to the condition in which a single basis kernel was used (**Figure 5**).

**Figure 5.**
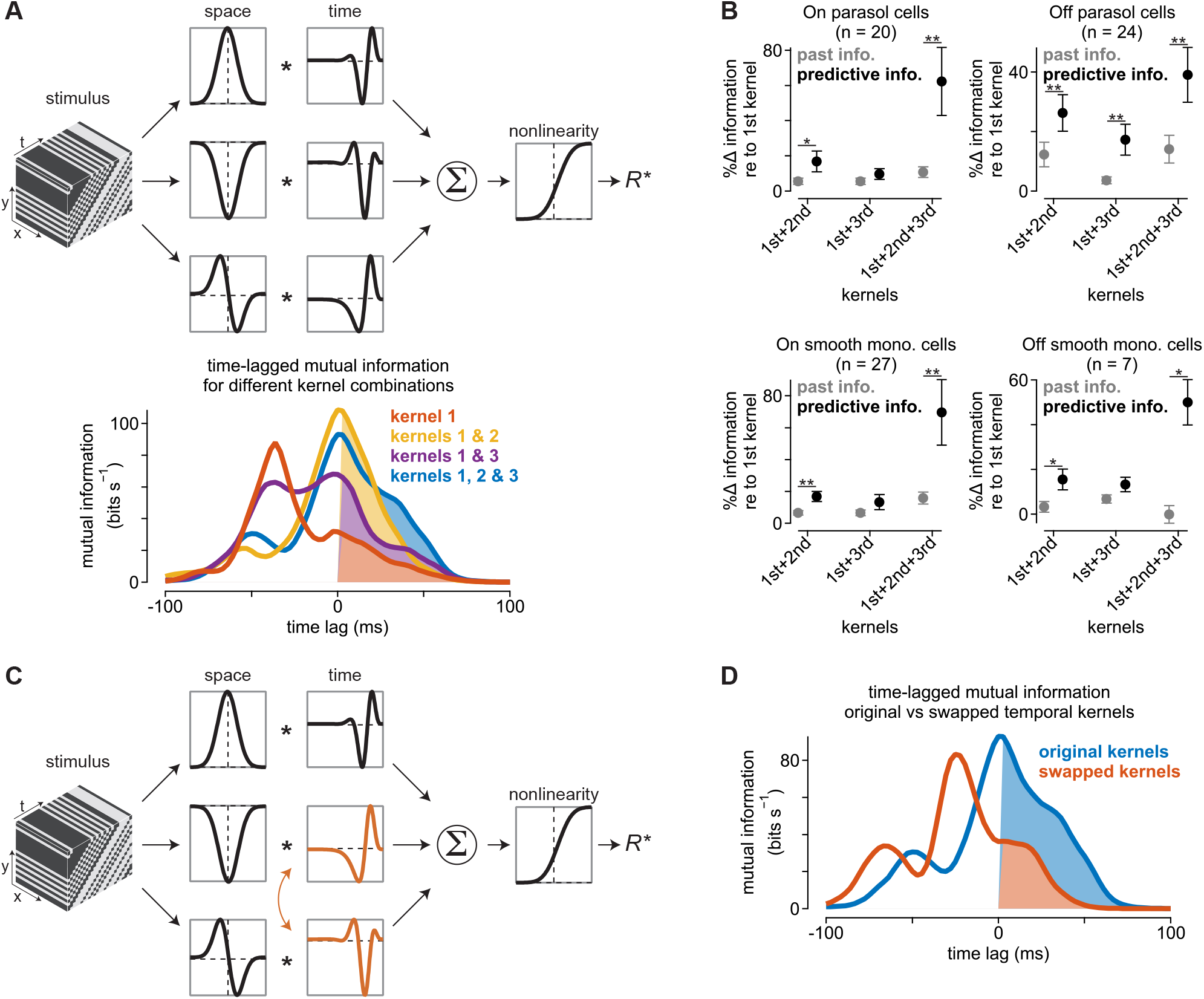
Additional receptive-field kernels improve predictive motion encoding. (A) *Top*, Spatiotemporal filtering of the stimulus in the model was performed using three space-time separable kernels estimated for each cell. The outputs of these spatiotemporal filters were summed and passed through a shared nonlinearity to produce a low-rank estimate of the neural response (*R*^∗^). Different weight combinations were used to estimate the contribution of the kernels to encoding. *Bottom*, Time-lagged mutual information curves for the different kernel combinations in the model shown. Shaded regions indicate the predictive information. The greatest predictive information was observed when all three spatiotemporal kernels were combined. (B) Population analysis showing the change in encoded information for different kernel combinations relative to the use of the dominant kernel alone. Results are shown for On parasol (n = 20), Off parasol (n = 24), On smooth monostratified (n = 27), and Off smooth monostratified (n = 7) cells. Inclusion of the second and third kernels in the low-rank receptive field estimate generally improved information encoding with the greatest improvement occurring for predictive information. Circles and error bars indicate mean ± SEM. Single asterisks indicate p-values <0.05 and double asterisks indicate p-values <0.005 (Wilcoxon signed rank test). (C) Model identical to that in (A) except that the temporal components of second and third kernels were swapped. (D) Mutual information curves for the original kernels (*blue*) from the model in (A) versus the swapped kernels from the model in (C). Shaded regions indicate the predictive information. Swapping the temporal kernel components decreased the encoding of predictive information.

We found that the additional kernels showed distinct effects on the encoding of past versus predictive motion information (**Figure 5**B). The additional kernels either weakly increased or had no effect on the encoded past information relative to the condition in which only the dominant kernel was used. These additional kernels did, however, increase predictive motion encoding with predictive information increasing by an average of >35% with the addition of the second and third kernels. These results indicate that the additional receptive field kernels improve motion encoding in these ganglion cells—particularly predictive motion encoding.

To determine whether the spatial and temporal components of the kernel basis were interchangeable, we swapped the temporal components of the second and third bases and recomputed the time-lagged mutual information between the stimulus and the model output (**Figure 5**C). The mutual information curves for the original and swapped kernels were distinct, with the original kernel producing a larger amount of predictive information than the swapped kernel (**Figure 5**D, *shaded regions*). These results indicate that the particular combination of spatial and temporal features present in the measured kernels are important for predictive encoding.

The second kernel recovered in On smooth monostratified and On parasol cells was suppressive, as positive projections along this kernel decreased the ganglion cell spike outputs (**Figure 3**). This suppressive (second) kernel further showed a large (~20 ms) delay in the peak response relative to the first kernel, and this delay likely explains the contribution of the kernel to predictive encoding for pairwise correlations. To test this hypothesis, we recomputed the model output after shifting the second kernel in time and recalculated the mutual information between the model output and the stimulus (**Figure 6**). The peaks of the shifted kernels are shown relative to the peak of the first kernel to illustrate the effects of the time delay on motion encoding. Indeed, information encoding varied as a function of this time shift—predictive information was highest when the second kernel peaked ~20-30 ms after the first kernel. Thus, the time delay between the suppressive kernel and the first kernel improved information encoding.

**Figure 6.**
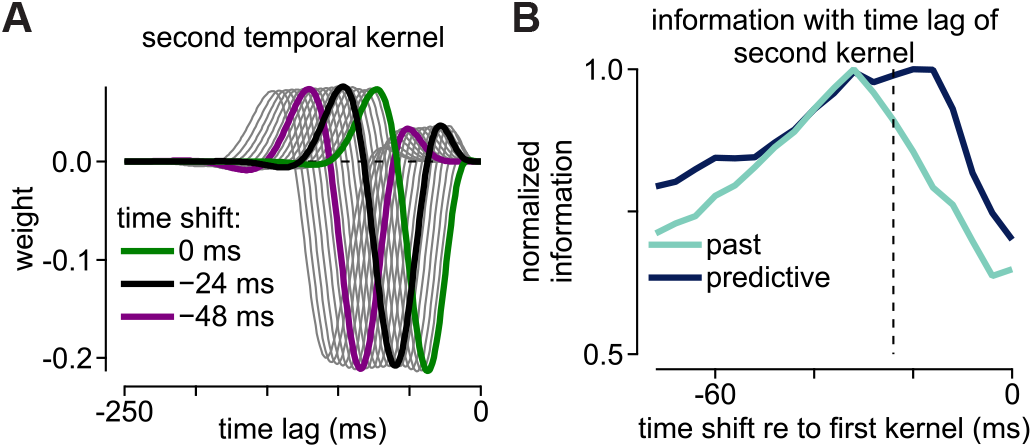
Relationship between the time lag of the second kernel and information encoding. (A) The relationship between the time lag of the second kernel and information encoding was investigated by shifting the second kernel to adjust the time at which this kernel reached a minimum value relative to the peak of the first kernel. (B) Normalized information for the model as a function of time shift in the second temporal kernel relative to the first kernel. The encoded predictive information peaked at time lags near the time lag of the estimated second kernel (dashed line).

Why would this time delay improve predictive encoding? The polarity of this kernel was opposite to that of the dominant kernel, suggesting that it suppressed spiking in the cell following a time delay. Delayed suppression of the spike response suppresses subsequent spiking and causes the peak spike response to occur earlier in time (Johnston and Lagnado, 2015; Berry et al., 1999; Leonardo and Meister, 2013; Schwartz et al., 2007). Our findings here further indicate that the timing of the temporal delay is critical to this mechanism. Short delays likely suppressed many of the faster, more informative spikes, while long delays were likely ineffective at speeding the peak spike response (see Discussion).

### Nonlinear subunits produce derivative receptive field modes

The spatial profiles of many cells in the visual cortex resemble the first derivative of a Gaussian function, similar to the structure we observed in our ganglion cell recordings (**Figure 2**). However, it was unclear how components of the retinal circuit contribute to this spatial structure. To investigate this question, we developed a subunit model of the bipolar cells providing inputs to parasol and smooth monostratified ganglion cells. Bipolar cell spatial properties were determined from direct measurements of excitatory synaptic currents from ganglion cells (Manookin et al., 2018; Appleby and Manookin, 2020; Liu et al., 2021). Following spatiotemporal filtering of the stimulus in the model bipolar cells, the input from each bipolar cell was passed through an input-output function that was either linear or nonlinear, after which the outputs were pooled at the level of the model ganglion cell. The model ganglion cell response was then used to extract the receptive-field structures as in **Figure 2**.

The first extracted filter for the linear subunit model showed a Gaussian spatial structure and biphasic temporal structure that was typical of the spike triggered average from a parasol or smooth monostratified ganglion cell (**Figure 7**A). However, the additional filters extracted from the analysis were dominated by noise and lacked clear spatiotemporal structure. This result indicated that the presence of receptive field subunits alone was not sufficient to produce the additional kernels that were present in our neural recordings and that contributed to predictive motion encoding.

**Figure 7.**
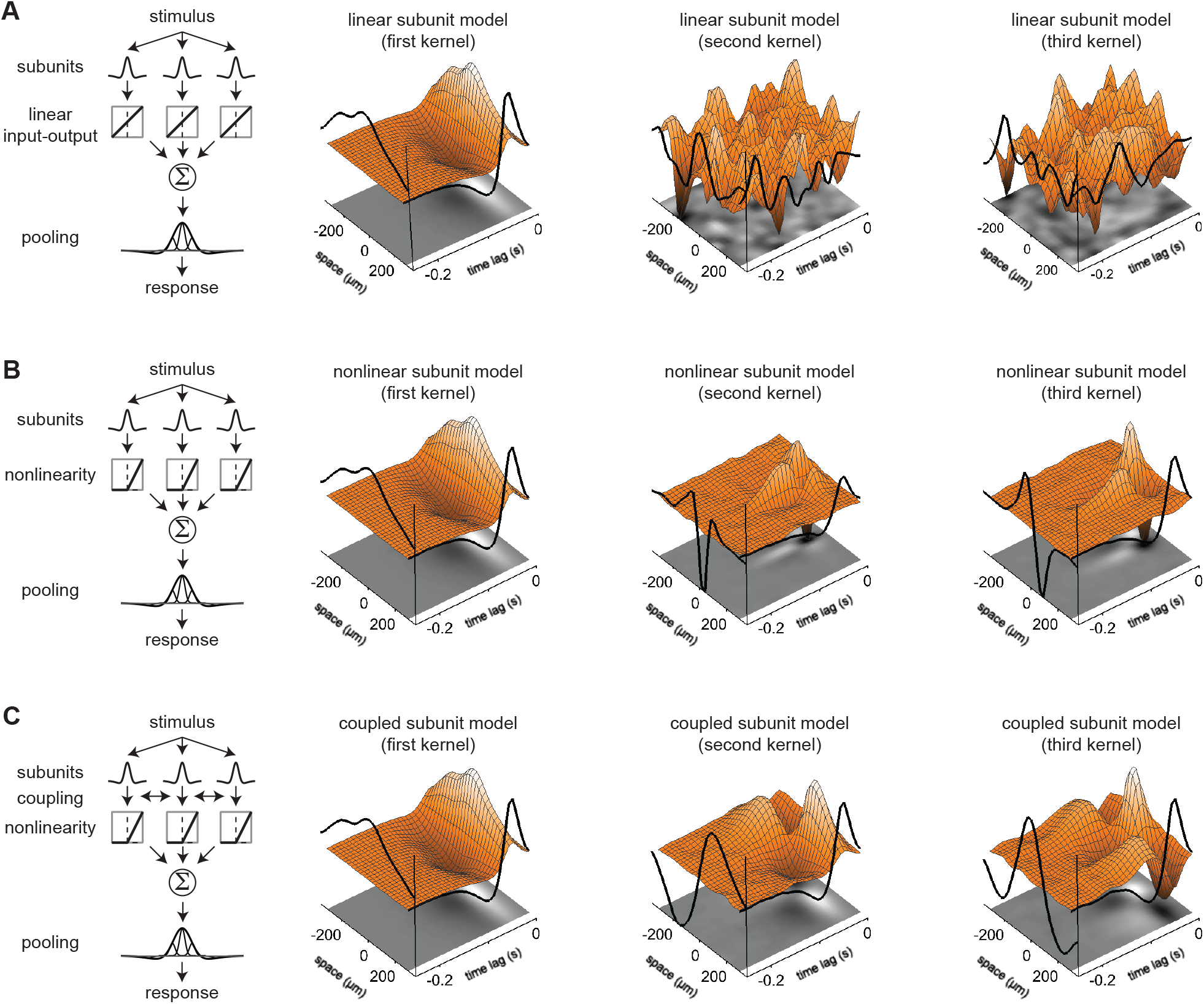
Nonlinear subunits sufficient to produce Gaussian derivative spatial kernels. (A) First three spatiotemporal kernels recovered for a subunit model in which the input-output relationship of the model bipolar cells was linear. The stimulus was 20 adjacent bars and the contrast of each bar was drawn pseudo-randomly from a Gaussian distribution in each time bin. The stimulus was filtered through the spatiotemporal receptive field of each model bipolar cell. The output of the filtering stage was then passed through the bipolar cell input-output function, after which the subunit signals were pooled and summed at the level of the model ganglion cell. The first filter showed a classical Gaussian spatial profile, but the second and third filters were dominated by noise and lacked any discernible spatiotemporal structure. (B) Spatiotemporal kernels for a model identical to that in (A) except that the input-output function for the model bipolar cell subunits was a piecewise nonlinearity (i.e., ReLU). The second kernel showed a spatial profile similar to the first derivative of a Gaussian function as was observed in the direct ganglion cell measurements. (C) Kernels for a subunit model identical to (B) except that electrical coupling was included between bipolar cell subunits. The derivative kernel was also observed, but was slightly smoother and more diffuse than that observed in (B).

Which properties of the retinal circuit could give rise to these additional receptive-field structures? The diffuse bipolar cells that provide synaptic input to parasol and smooth monostratified ganglion cells show strongly nonlinear relationships between their inputs and their synaptic outputs (Turner and Rieke, 2016; Manookin et al., 2018). Thus, we tested whether this nonlinear processing contributed to the additional receptive field modes. This nonlinear subunit model was identical to the linear subunit model except for the non-linear subunit output. Indeed, including a nonlinearity at the model bipolar cell output resulted in additional receptive-field kernels that resembled the first derivative of a Gaussian function, similar to what was observed in our direct ganglion cell recordings (**Figure 7**B).

The kernels computed for the nonlinear subunit model showed a derivative structure, but the spatial extent of the structures were much smaller than those observed in our direct recordings and in the model in which the subunits were coupled (**Figure 2**, **Figure 4**, **Figure 7**C). A possible explanation of this is that the kernel structures in **Figure 7**B are dominated by only a few of the subunits. To test this, we modified the model to increase the integration area in the model ganglion cell and recomputed the kernel estimates (**Figure 8**). For the nonlinear subunit model that lacked subunit coupling, tripling the diameter of the ganglion cell receptive field did not dramatically affect the size of the recovered kernels (**Figure 8**A). Instead, the coupled subunit model with both coupling between the subunits and a nonlinearity at their outputs best reproduced the receptive field kernels measured from the direct recordings. These findings indicate that both nonlinear input-output functions of bipolar cells and electrical coupling are necessary to explain both the shape and the extent of the derivative spatial filters observed in parasol and smooth monostratified ganglion cells.

**Figure 8.**
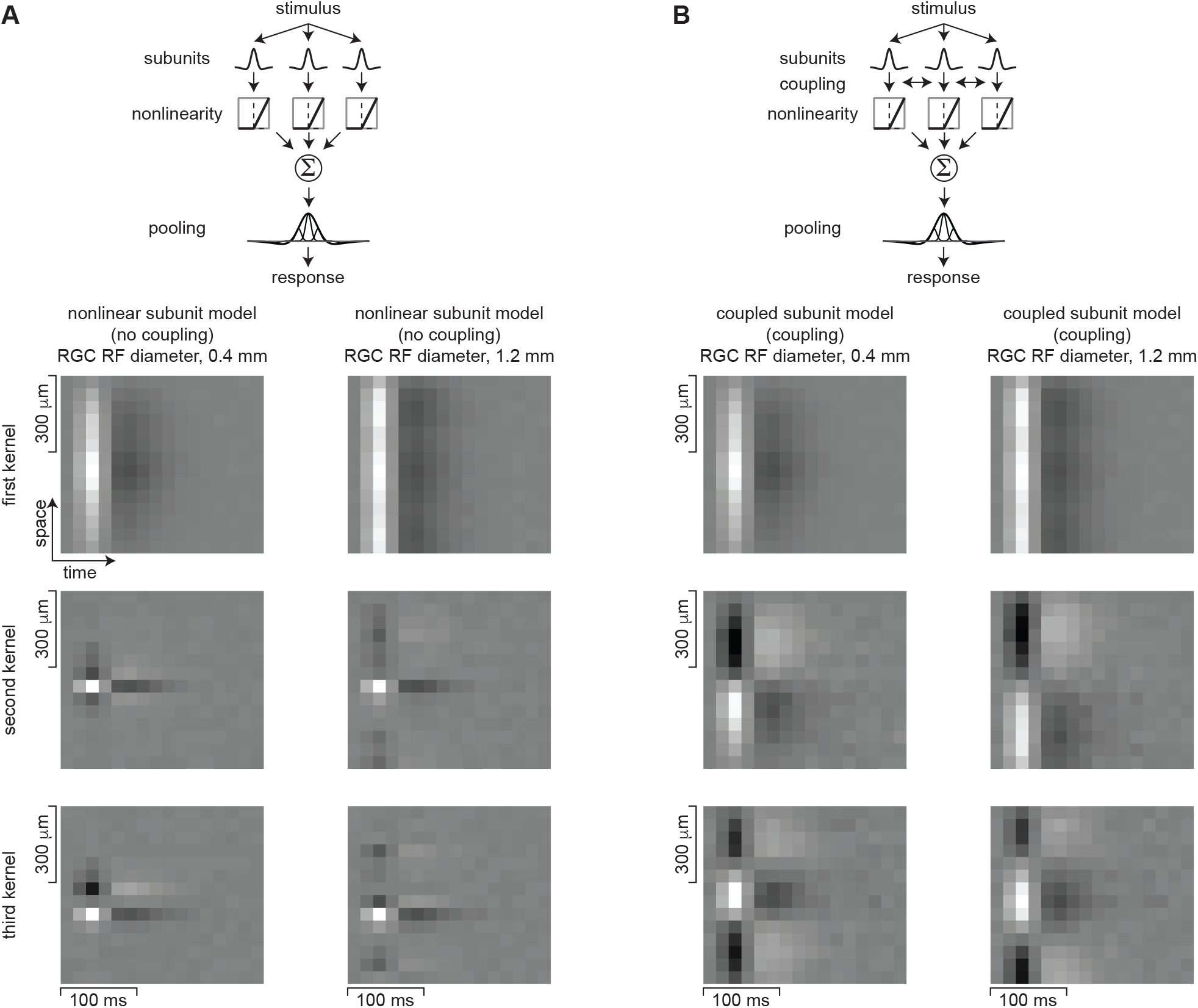
Electrical coupling between nonlinear subunits needed to explain spatial extent of receptive field kernels. (A) The first three spatiotemporal kernels computed for the nonlinear subunit model. Results are shown for ganglion cell receptive field diameters of 0.4 mm (*left column*) and 1.2 mm (*right column*). The recovered kernels for the two models were small relative to the ganglion cell receptive field diameter, suggesting that they primarily arose from only a small number of subunits. (B) Results for the coupled subunit model, which was identical to that in (A) except that the subunits were coupled. The recovered spatial structures increased with the ganglion cell receptive field size and more closely resembled the measured receptive field kernels in parasol and smooth monostratified ganglion cells. This indicates that subunit coupling could also contribute to the derivative receptive field structures.

### Sparse spatial integration improves encoding of predictive motion information

With few documented exceptions (Manookin et al., 2015), the receptive field centers of cells in the primate retina are well described by a single Gaussian function. Smooth monostratified ganglion cells constitute a clear exception to this rule—these cells show spotty receptive fields that sparsely sample visual space, but the potential contributions of this sparsity to predictive encoding is not understood. (Rhoades et al., 2019). To determine how sparse spatial sampling contributed to motion encoding, we extended our subunit model so that the subunit outputs were either pooled using a Gaussian receptive field or a sparse receptive field (**Figure 9**). This sparse spatial receptive field was directly measured from an On-type smooth monostratified cell using an uncorrelated spatiotemporal noise stimulus (**Figure 9**B, *left*). Other than the spatial pooling component, the two models were identical. We calculated the mutual information between the model spike output and stimulus for both models, and past versus predictive information were measured.

**Figure 9.**
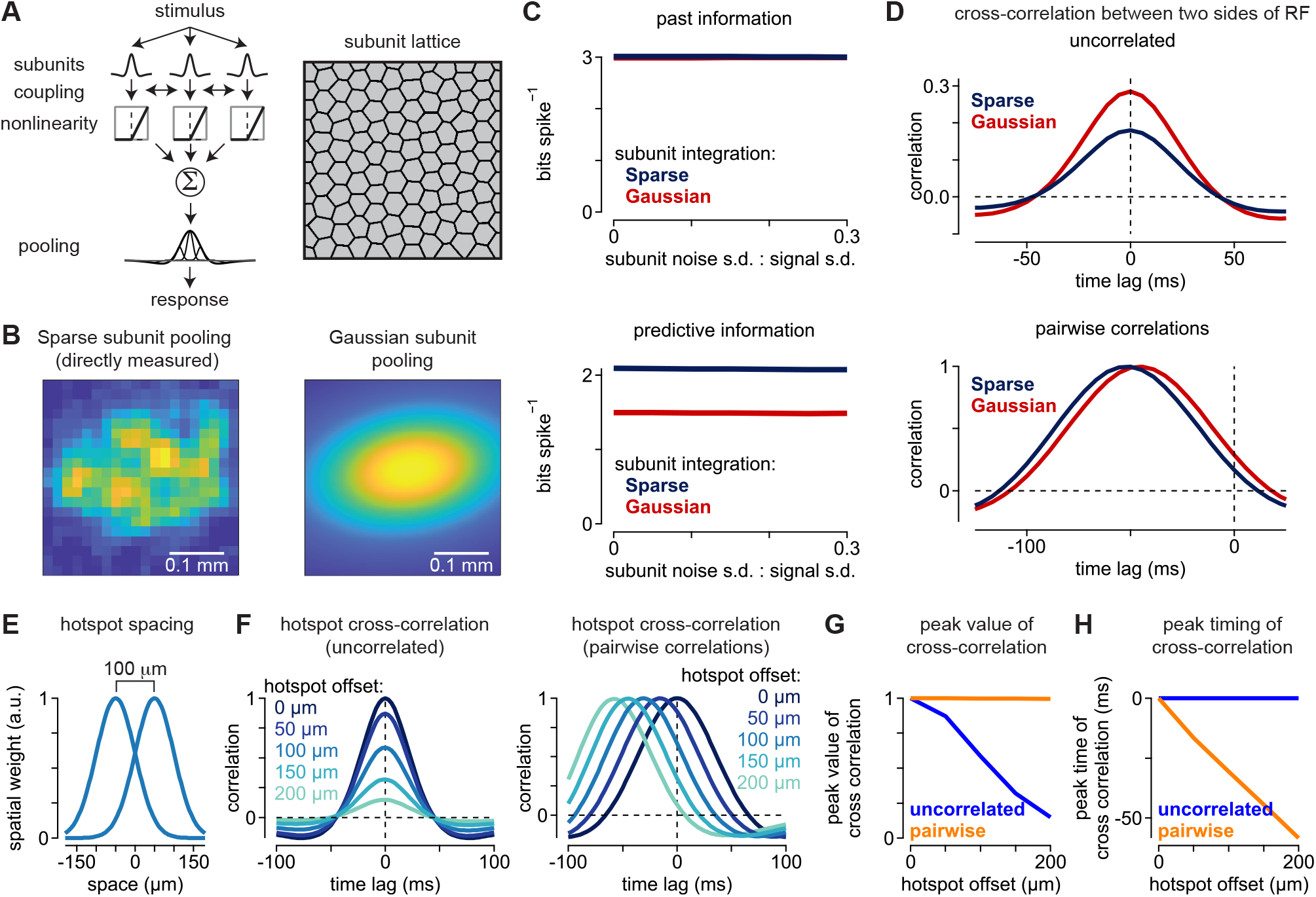
Sparse pooling of receptive-field subunits increases predictive motion encoding. (A) Subunit model organization. The stimulus was filtered by the spatiotemporal receptive field of each model bipolar subunit. A portion of the response in each bipolar cell was shared with neighboring bipolar cells via electrical coupling after which the model bipolar currents were passed through a piecewise nonlinearity. Pooling of these rectified signals then occurred at the level of the model ganglion cell. (B) Pooling of the signals from model subunits occurred either using a spatial receptive field profile that was directly measured in a smooth monostratified ganglion cell (*left*) or using a two-dimensional Gaussian fit to the receptive field (*right*). Scale bars, 0.1 mm. (C) *Top*, Past information encoded in bits spike^−1^ as a function of the amount of additive noise in the individual subunits prior to coupling and the nonlinear output. Noise is shown as the ratio between the noise standard deviation and the signal standard deviation. Gaussian pooling and sparse pooling produced similar amounts of past information encoding. *Bottom*, Encoded predictive information for the two models. Sparse pooling of bipolar subunits produced higher encoding of predictive information than Gaussian pooling across noise levels. (D) Cross-correlation between the left and right halves of the receptive field for an uncorrelated stimulus (*top*) and a stimulus containing pairwise spatiotemporal correlations (*bottom*). The uncorrelated stimulus produced lower correlation values for the sparse sampling versus Gaussian sampling of the subunits (*top*). For the pairwise correlations, peak correlation values were similar, but sparse sampling produced a larger shift in the temporal lag between the two sides of the receptive field, consistent with the higher degree of predictive encoding in (C). (E) Simplified model of spatial sparsity in which the receptive field was comprised of two identical Gaussian hotspots that varied only in their spatial offsets (*σ*, 50 *μm*; offset, 0-200 *μm*). The hotspots independently integrated the subunit outputs. (F) Cross-correlation between the two hotspots for the uncorrelated (*left*) and pairwise correlation stimuli (*right*). As separation between the hotspots increased, the correlation decreased for the uncorrelated stimulus, but remained unchanged for the motion stimulus. However, the peak of the cross-correlation occurred earlier for the motion stimulus as separation increased. (G) Correlation as a function of hotspot offset for the uncorrelated stimulus (*blue*) and pairwise correlations (*orange*). (H) Timing of the peak of the cross-correlation as a function of hotspot offset for the uncorrelated stimulus (*blue*) and pairwise correlations (*orange*).

The sparse pooling and Gaussian pooling models showed distinct encodings of past versus predictive motion information. Encoded past information was similar for the models (**Figure 9**C). However, a different pattern was observed for predictive encoding—sparse pooling of the subunit outputs produced a higher encoding of predictive information relative to Gaussian pooling of the same subunit outputs. This result indicates that sparse spatial sampling biases predictive information during neural encoding.

Mechanisms that speed the neural response tend to increase the predictive encoding of motion (Berry et al., 1999; Schwartz et al., 2007; Leonardo and Meister, 2013; Johnston and Lagnado, 2015; Liu et al., 2021). Similarly, sparse spatial sampling could also cause spiking to occur earlier and, thus, increase predictive encoding. The sparsity of smooth monostratified cell receptive fields is characterized by areas of sensitivity concentrated at the margins of the receptive field and a relative lack of sensitivity in the center [see Figure 3 of (Rhoades et al., 2019)]. Thus, a sparse sampling may cause a cell to respond earlier as a moving object encroaches upon the edge of the receptive field than if the sampling were Gaussian with the highest sensitivity regions concentrated toward the receptive field center.

To test this idea, we computed the cross-correlation between the two halves of the model ganglion cell receptive field during the stimulus with pairwise motion correlations. If the cell were responding earlier, then the peak of the cross-correlation would be shifted to earlier time points. Indeed, the cross-correlation peaked earlier for the model with sparse spatial sampling than for the Gaussian sampling model (**Figure 9**D). This result indicates that the sparse integration model showed higher levels of predictive motion encoding relative to the Gaussian model, in part, because the sparsity of the receptive field caused motion responses to occur earlier in time (see Discussion).

### Neural adaptation enhances predictive encoding

The Gaussian derivative spatial kernels that we observed in parasol and smooth monostratified cells increase a cell’s sensitivity to changes occurring across space. These cells also show strongly biphasic temporal kernels, which further increase their ability to detect changes in time (Rhoades et al., 2019). Together these receptive field components, largely inherited from their presynaptic inputs, increase a cell’s ability to detect the changes in space and time that occur during visual motion (Kuo et al., 2016; Manookin et al., 2018). Neural adaptation is an additional mechanism that increases a cell’s ability to detect changes in their inputs (Fairhall et al., 2001; Smirnakis et al., 1997).

Adaptation adjusts a cell’s output to match the statistics of the incoming stimulus, which increases the cell’s sensitivity to changes in the stimulus. Indeed, adaptation was proposed as the principal mechanism for predicting translational motion in the salamander retina (Berry et al., 1999; Schwartz et al., 2007; Leonardo and Meister, 2013). These previous studies examined the contribution of the gain control mechanism in the context of transient motion into and out of the receptivefield center and only to pairwise spatiotemporal correlations. Thus, the potential contribution of gain control to predictive encoding for continuous motion and triplet spatiotemporal correlations has not been carefully studied.

To investigate whether adaptation contributes to predictive encoding for diverging and converging spatiotemporal correlations, we developed a computational model of smooth monostratified cells that included this mechanism. We estimated the temporal filtering and adaptation properties of bipolar cell inputs and spike outputs of a ganglion cell by recording excitatory synaptic currents or spike responses to a spatially uniform spot presented over the cell’s receptive field (**Figure S3**). The contrast of the spot was drawn randomly from a Gaussian distribution in each stimulus period (mean, 0.0; standard deviation, 0.3). The data were then analyzed using a generalized linear model (GLM) (Pillow et al., 2008; Paninski, 2004; Truccolo et al., 2005). In addition to modeling the temporal filtering properties of a cell, this model framework accounts for the modulation of neural output based on the recent history of neural responses (an adaptation filter). For this reason, generalized linear models have been useful in modeling adaptation in neurons (Latimer and Fairhall, 2020; Weber and Pillow, 2017; Latimer et al., 2019; Mease et al., 2013).

The generalized linear model comprised three processing stages: 1) a temporal kernel that filtered the incoming stimulus, 2) a point nonlinearity that mapped the output of the temporal filtering stage to a neural output (spikes or conductance), and 3) an adaptation filter that provided feedback to the output of the temporal filtering stage based on the recent neural output. This final stage behaved similarly to gain control mechanisms that suppress neural responses following strong outputs (Latimer and Fairhall, 2020; Weber and Pillow, 2017). To measure the time course of adaptive feedback, we fit the adaptation filters with a single exponential. The adaptation decayed rapidly for both spiking and excitatory synaptic currents, indicating that this feed-back suppressed neural responses on relatively short time frames (decay time constant: spiking, 5.9 ms; excitatory currents, 5.0 ms; **Figure S3**D).

To determine whether adaptation influenced predictive encoding for pairwise and triplet spatiotemporal correlations, we incorporated the empirically determined filters into our circuit model. The adaptation filters were implemented at one of two sites in the model—either at the bipolar cell or ganglion cell outputs (**Figure 10**). The output of this filter was normalized to the same standard deviation as the output of the temporal filter so that the contribution of adaptation could be properly quantified. We tested whether the magnitude of adaptation affected predictive encoding by varying the weight of the adaptation filter (weight, 0-1) with a weight of zero corresponding to a model lacking adaptation. We further tested for interactions between adaptation and neural sparsity by varying the fraction of time bins in which spiking occurred.

**Figure 10.**
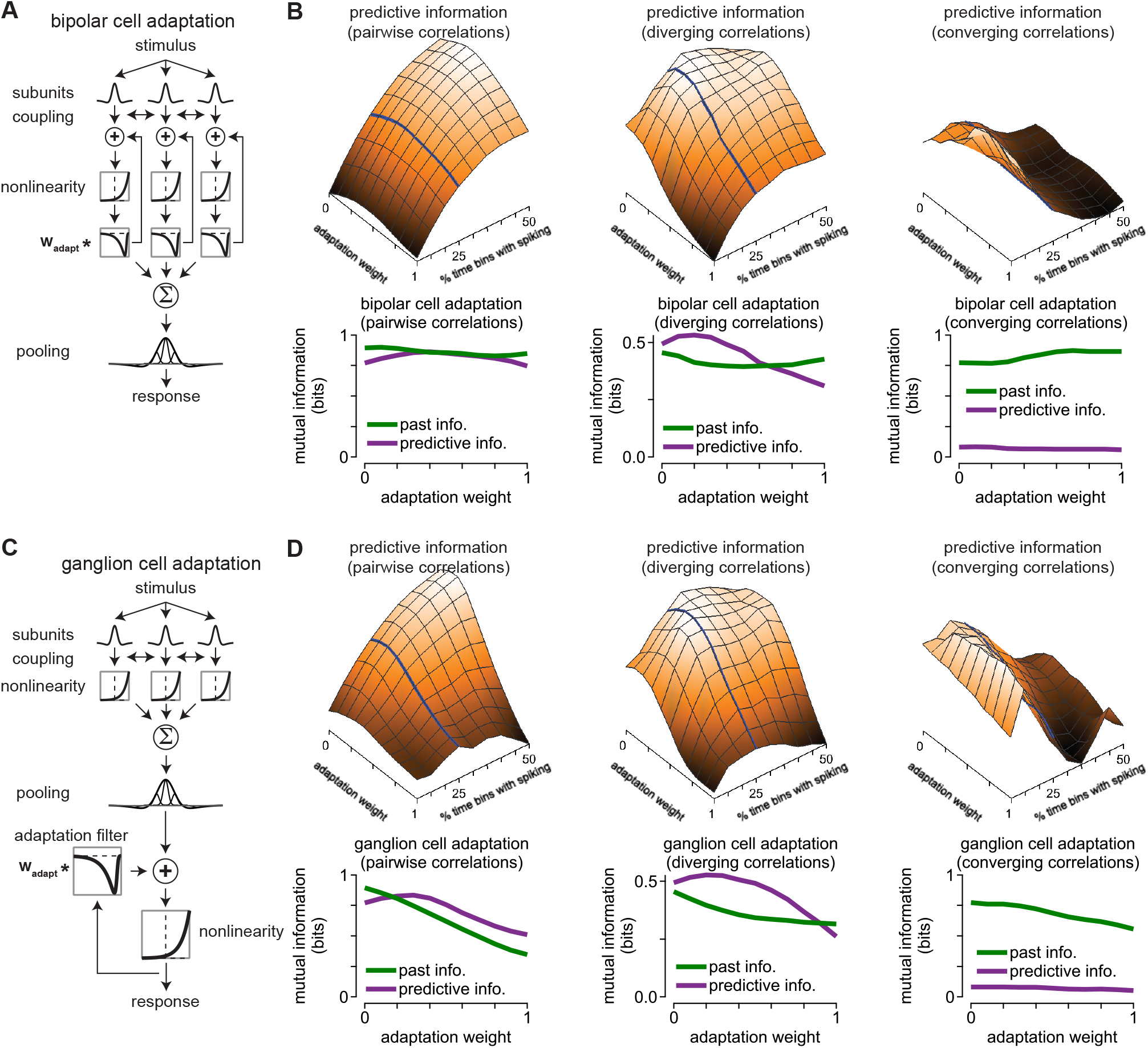
Moderate adaptation improves predictive encoding for spatiotemporal correlations. (A) Organization of the bipolar cell adaptation model. An adaptation filter was applied to model signals based on the recent output of each model bipolar cell subunit—stronger outputs resulted in greater suppression of model signals prior to the output nonlinearity. (B) Contribution of bipolar cell adaptation to encoding of past and future (predictive) information. *Top*, Surface showing the encoded predictive information as a function of the adaptation weight and temporal sparsity (percentage of time bins containing a spike). Surfaces are shown for pairwise (*left*), diverging (*middle*), and converging (*right*) spatiotemporal correlations. *Bottom*, Past and predictive information as a function of adaptation weight. Curves are shown for a temporal sparsity value of 25% (*top*, solid line). (C) Organization of a model in which adaptation occurs at the ganglion cell level. (D) Information curves as in (B) for the ganglion cell adaptation model. Predictive information encoding peaked for adaptation weights of 0.2-0.4 and decreased at higher values.

These model simulations indicated that moderate gain control was beneficial to predictive motion encoding (**Figure 10**B, D). The computed predictive information peaked near adaptation weights of ~0.2-0.4 and decreased at lower and higher values. This trend was observed for the models with both sparser and denser temporal coding (**Figure 10**B, D, *top row*), and it was also true for the models in which adaptation occurred at either the bipolar cell or ganglion cell outputs. Observing these results across a range of model conditions highlights the benefit of moderate adaptation in predictive encoding. Thus, consistent with previous findings, moderate adaptation increased predictive motion encoding (Berry et al., 1999; Schwartz et al., 2007; Leonardo and Meister, 2013).

Mechanisms such as adaptation that speed the neural response are generally considered beneficial to predictive encoding (Berry et al., 1999; Schwartz et al., 2007; Leonardo and Meister, 2013). Similar to delayed inhibition (**Figure 6**), following strong neural responses, adaptation mechanisms provide negative feed-back, which decreases subsequent responses (Kim and Rieke, 2001; Fairhall et al., 2001; Baccus and Meister, 2002). This effectively makes responses peak earlier and increases the amount of predictive information in the neural output. However, our results indicate that adaptation is advantageous only within a fairly limited range—when the magnitude of adaptation exceeded ~40% of the spatiotemporal filter output, predictive encoding was suppressed.

This range in which adaptation supports predictive encoding may reflect a tradeoff between speeding the neural response by removing spikes (moderate adaptation) and removing informative spikes that degrade information encoding (strong adaptation). For example, moderate adaptation (weight, 0.3) increased the mutual information at positive time lags relative to the unadapted condition, resulting in an increase in predictive information (**Figure S4**). However, the excessive suppression of spiking caused by strong adaptation caused a net decrease in information at positive time lags relative to the unadapted condition.

## DISCUSSION

A central pursuit of computational and systems neuroscience is to understand the relationship between stimuli in the external environment and neural responses. Here, we studied how properties of the retinal circuit contribute to motion encoding in primates. We found that several circuit properties collectively improved the ability of parasol and smooth monostratified ganglion cells to encode information about visual motion. This improvement was particularly evident for predictive motion encoding—the ability of the cell to convey information about the future trajectory of moving objects (**Figure 2**, **Figure 4**, **Figure 5**, **Figure 6**, **Figure 9**). Non-linear mechanisms such as the rectified synaptic release from bipolar cells and adaptation further enhanced predictive motion encoding (**Figure 7**, **Figure 10**, **Figure S4**). Thus, several properties of parasol and smooth monostratified ganglion cells support accurate estimation of trajectories of moving objects.

Several receptive-field properties that contribute to predictive motion encoding are strong candidates for contributing to predictive computations in other sensory regimes. For example, the spatiotemporal derivative kernel improved predictive motion estimation, and similar derivative kernels are found in both the visual and auditory regions of the cortex (DeAngelis et al., 1993a,b; De Valois and Cottaris, 1998; deCharms et al., 1998; Singer et al., 2018). Furthermore, delayed suppression of the neural response from adaptation mechanisms and synaptic inhibition are common features of neural circuits throughout the brain. This suppression contributes to prediction by speeding neural responses and thus overcoming some of the temporal delays inherent in neural processing (Berry et al., 1999; Schwartz et al., 2007; Johnston and Lagnado, 2015). Finally, sparse signal integration is another mechanism, identified here, that could contribute to predictive computations in other neural systems (**Figure 9**). This mechanism is discussed further in the following text.

### Linear receptive field properties improve motion estimation

The encoding of correlations is at the core of the predictive computation. In principle, two points within the receptive field that are correlated with each other can participate in predictive encoding if the activity in one point at a particular time predicts the activity in the second point at a later time. Furthermore, this contribution to predictive encoding would occur even if the relationship between the points is linear. Our previous work focused on the contribution of two circuit properties to predictive motion encoding—electrical coupling and the bipolar cell synaptic output (Liu et al., 2021). The results presented here indicate that other receptive field properties also contribute to this computation.

The spatial receptive fields of parasol and smooth monostratified ganglion cells consistently showed a spatial kernel resembling the derivative of a Gaussian function (**Figure 2**). This structure’s importance lies in the adjacent On and Off subregions within the receptive field. The balanced weighting of these regions means that the output of this kernel will be weak or absent for stimuli that do not vary in intensity. However, responses will be strong for stimuli that vary in their intensity, such as when the edge of an object moves through the receptive field. Indeed, this spatial receptive field profile is common in cortical cells that contribute to motion processing (Adelson and Bergen, 1985; Emerson et al., 1992; Reid et al., 1987, 1991; Rust et al., 2005).

We treated this derivative spatial structure as a linear operator and assessed its contribution to motion encoding (**Figure 5**), but this structure can also arise as a property of nonlinear signaling in bipolar cells (**Figure 7**). This bipolar cell origin is key to understanding how stimuli will exercise the derivative spatial structure. While the derivative structures that we measured were oriented along the long axis of the bars that were presented, the orientation of this receptive field structure should be stimulus dependent. This pliancy differs from the properties of motion sensitive neurons in the visual cortex that show static receptive field orientations (Adelson and Bergen, 1985; Emerson et al., 1992; Reid et al., 1987, 1991; Rust et al., 2005). Thus, the representation of visual motion in parasol and smooth monostratified ganglion cells is simultaneously less selective and more flexible in its orientation than that found in downstream visual areas.

### Sparse spatial sampling improves predictive encoding

The spatial component of smooth monostratified ganglion cell receptive fields shows sparse sampling relative to many other mammalian ganglion cell types, but the functional implications of this sparsity are not known (Rhoades et al., 2019). We asked whether sparse sampling contributes to motion encoding by comparing two models that differed only in their spatial sampling—a uniform Gaussian sampling and a sparse sampling taken from direct receptive-field measurements (**Figure 9**). Indeed, past and predictive information encoding differed between these two models with sparse sampling encoding more predictive information than Gaussian sampling.

Sparse spatial sampling appears to benefit predictive encoding, at least in part, by causing a cell to respond earlier when a moving object encroaches on the edge of the receptive field than for a smooth receptive field (**Figure 9**D). Indeed, speeding of response kinetics is a critical component of motion anticipation in the salamander and fish retinas (Berry et al., 1999; Johnston and Lagnado, 2015; Leonardo and Meister, 2013; Schwartz et al., 2007). Smooth monostratified ganglion cells show sensitivity to stimuli falling only within limited regions within the receptive field, and these sensitive regions are separated by areas lacking sensitivity [(Rhoades et al., 2019); **Figure 9**B]. Moreover, these sensitive regions are typically found toward the edges of the receptive field. Thus, objects moving into the margins of the receptive field will tend to contact a sensitive region and evoke responses before contacting less sensitive regions. This causes responses to occur earlier than if the cell were sampling space with a Gaussian receptive field in which the strongest regions are located at the center.

This sparse sampling does sacrifice some spatial acuity as a cell will not respond to objects falling in certain regions of the receptive field. However, cortical neurons likely have access to signals from multiple ganglion cell types and these signals can then be combined in ways that allow cortical neurons to compute local motion signals on a finer spatial scale (Movshon and Newsome, 1996; Hubel and Wiesel, 1974). Thus, the spatial sampling in smooth monostratified cells may be sufficient for detecting visual motion and a benefit of this sparse sampling is that it promotes predictive encoding in a similar way to adaptation mechanisms—by biasing responses to moving objects at the edge of the receptive field (Berry et al., 1999).

### Delayed suppression improves predictive encoding

Several studies have highlighted adaptation (gain control) as the key mechanism contributing to predictive motion estimation in salamander retina (Berry et al., 1999; Leonardo and Meister, 2013; Schwartz et al., 2007). However, another study in the fish retina indicated that feedforward inhibition played the principal role in this computation (Johnston and Lagnado, 2015). Our findings here indicate that both mechanisms can work in concert to improve motion estimation.

The central idea is that these mechanisms work on different time scales to speed the neural response (**Figure 6**, **Figure 10**, **Figure S4**). Adaptation provides rapid feedback following strong spiking, which suppresses subsequent spiking and causes the peak spike response to occur earlier. This suppression occurs and decays rapidly and thus acts on relatively short time scales (decay time constant, 5.0-5.8 ms). Our results further indicate that the strength of this feedback must be properly tuned in order to be effective—strong adaptation suppressed informative spikes and degraded information encoding (**Figure 10**, **Figure S4**).

We also observed a suppressive kernel that showed a temporal delay relative to the dominant kernel. This kernel peaked approximately 20 ms after the dominant kernel and showed more sustained kinetics than adaptation (**Figure 2**). For the purposes of this study we do not claim that this kernel arises from amacrine cells, but the temporal delay and effects on predictive coding are qualitatively similar to those mediated by feedforward inhibition in the fish retina in that they both improved predictive motion estimation [**Figure 6**; (Johnston and Lagnado, 2015)]. Thus, this suppressive kernel and adaptation can modulate neural dynamics on different time scales and fine tune predictive motion information arising in the excitatory circuitry (Liu et al., 2021).

## ACKNOWLEDGEMENTS

We thank Shellee Cunnington for technical assistance. Tissue was provided by the Tissue Distribution Program at the Washington National Primate Research Center (WaNPRC; supported through NIH grant P51 OD-010425), and we thank the WaNPRC staff, particularly Chris English and Audrey Baldessari, for making these experiments possible. Chris Chen assisted in tissue preparation. This work was supported in part by grants from the NIH (NEI R01-EY027323 to M.B.M.; NEI R01-EY029247 to E.J. Chichilnisky, F.R., and M.B.M.; NEI R01-EY028542 to F.R.; NEI P30-EY001730 to the Vision Core), Research to Prevent Blindness Unrestricted Grant (to the University of Washington Department of Ophthalmology).

## AUTHOR CONTRIBUTIONS

Conceptualization, M.B.M.; Methodology, B.L., A.H., F.R., and M.B.M.; Software, M.B.M.; Formal Analysis, B.L. and M.B.M.; Investigation, M.B.M.; Resources, F.R. and M.B.M.; Data Curation, M.B.M.; Writing – Original Draft, M.B.M.; Writing – Review and Editing, B.L., A.H., F.R., and M.B.M.; Visualization, B.L and M.B.M.; Supervision, F.R. and M.B.M.; Project Administration, F.R. and M.B.M.; Funding Acquisition, F.R. and M.B.M.

## METHODS

Experiments were performed using an *in vitro*, pigment-epithelium attached preparation of the macaque monkey retina from three different macaque species of either sex (Macaca *fascicularis*, *mulatta*, and *nemestrina*). Tissues were obtained from terminally anesthetized animals that were made available through the Tissue Distribution Program of the National Primate Research Center at the University of Washington. All procedures were approved by the University of Washington Institutional Animal Care and Use Committee.

Recorded cells were located in the macular, mid-peripheral, or peripheral retina (2-8 mm, 10-30° foveal eccentricity). Data were acquired using a Multiclamp 700B amplifier (Molecular Devices), digitized using an ITC-18 analog-digital board (HEKA Instruments), and acquired using the Symphony data acquisition software (http://symphony-das.github.io). Other analyses of this dataset are published elsewhere (Liu et al., 2020, 2021).

### Visual stimuli

Visual stimuli were generated using the Stage software package (http://stage-vss.github.io) and displayed on a customized digital light projector (Appleby and Manookin, 2019, 2020). Stimuli were presented at medium to high photopic light levels with average L/M-cone photoisomerization rates (R*) of ~1.5 × 10^4^ – 5.0 × 10^5^ s^−1^.

### Receptive-field kernel estimation

Our goal was to describe the relationship between the stimulus (***s***) and a cell’s spike output (**r**) using three spatiotemporal kernels (***K***). The computing time required to run the algorithm made calculating more than three kernels for each cell computationally intractable. We estimated the kernels that maximized the average information conveyed by a single spike about the stimulus projected onto ***K*** (Sharpee et al., 2004; Williamson et al., 2015). First, the prior stimulus distribution was determined by projecting the stimulus onto a candidate kernel basis (*p*(***K***^⊤^***s***)) and the spike-triggered distribution was determined by projecting the stimuli that elicited spiking onto this basis (*p*(***K***^⊤^***s***|*spike*)). The single-spike information (*I_spike_*) was then determined by calculating the separation between these distributions using the Kullback-Leibler divergence (Williamson et al., 2015).

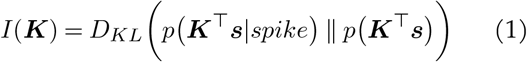

where *D_KL_* is the Kullback-Leibler divergence and *p*(***K***^⊤^***s***) and *p*(***K***^⊤^***s***|*spike*) are the raw and spike-triggered stimulus distributions projected onto ***K***.

The nonlinear relationship between the stimulus projection onto the kernel basis (***K***) and the spike rate of the cell was determined using an exponential mapping between the stimulus projection onto the basis and the spike output of the cell (**r**). To aid in fitting, the nonlinearity was parameterized using radial basis functions (*ϕ*, Equation 2).

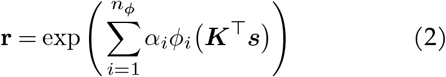

where *α* are the linear weights on the radial basis functions (Williamson et al., 2015).

### Estimation of 1D and 2D nonlinearities

The shared nonlinearity between the kernels was determined by computing the spiking probability conditioned on the stimulus projection onto the individual kernels. The individual kernel nonlinearities were then determined by computing the average spike probability along each kernel projection axis.

The two-dimensional nonlinearity representing the condition in which each kernel had a separate nonlinearity was then determined by taking the outer product of the average kernel nonlinearities (Equation 3), and then scaling this nonlinearity such that the total spike probability matched that of the shared nonlinearity.

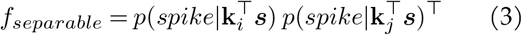

where **k**_*i*_ and **k**_*j*_ are the *i*th and *j*th spatiotemporal kernels.

### Spatial kernel modeling

We modeled the spatial component of the kernel estimates as either a Gaussian function (Equation 4) or the derivative of a Gaussian (Equation 5).

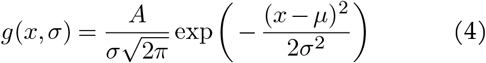

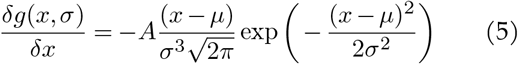

where *μ* is the spatial offset of the receptive-field center in microns, *σ* is the standard deviation in microns, and *A* scales the amplitude of the resulting function.

The spatial component of each kernel was fit with both functions and goodness-of-fit was determined by calculating the Pearson correlation (*r*^2^) between the fit and the raw data (see **Figure 2**).

### Generalized linear model

We used a generalized linear model framework to represent temporal filtering and adaptation at either the bipolar cell synaptic output or the ganglion cell spike output. The time-varying neural response (**r**_*t*_) was modeled as a nonlinear function (*f*) of the projection of the temporal kernel (**k**) onto the stimulus (**s**_*t*_) summed with the projection of a filter that captures the history-dependence of the neural response (**h**) onto the history of neural responses (**y**_*history,t*_).

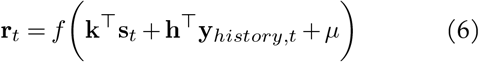

where the rows of **s**_*t*_ are time samples and the columns are the stimulus vector in the 500 ms preceding time *t*. Similarly, the rows of the response history matrix (**y**_*history,t*_) are time samples and the columns are the neural responses the 100 ms preceding time *t*. The scalar variable *μ* represents the maintained neural response.

For the excitatory synaptic signals measured in voltage-clamp, the synaptic conductance was used in place of the current so that positive values correspond to increases in excitatory input. Conductance (*g*) was calculated as the ratio of the synaptic current (*I*) and the driving force:

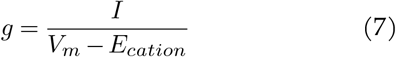

where *V_m_* is the membrane potential (−70 mV) and *E_cation_* is the cation reversal potential (0 mV).

The kernel coefficients 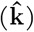 were then estimated using ridge regression:

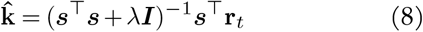

where ***I*** is the identity matrix and *λ* is the ridge parameter.

### Computational model

We created a model of the diffuse bipolar cells that provide excitatory synaptic input to parasol and smooth monostratified ganglion cells. A lattice of model bipolar cells was created with a mean spacing of 32 microns (Boycott and Wässle, 1991; Tsukamoto and Omi, 2015, 2016). The spatiotemporal filtering and output nonlinearities for the bipolar cells were determined by direct measurements (Appleby and Manookin, 2020; Manookin et al., 2018). The spatiotemporal receptive field of each bipolar cell (***F***_*i*_) was generated from the outer product of the Gaussian spatial component and a biphasic temporal component and the linear response of each bipolar cell (**r**_*i*_) was determined by projecting the stimulus onto its receptive field.

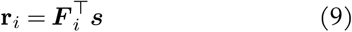

Noise in the bipolar responses was simulated by adding Poisson fluctuations to the resulting bipolar cell responses and coupling between the cells was applied based on our direct measurements (Manookin et al., 2018). The response of each bipolar cell following coupling was determined by adding the change due to coupling to the response prior to coupling (*R*_0_; Equation 10).

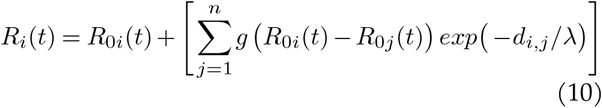

where *g* is the coupling gain or portion of the response shared between bipolar cells, *λ* is the coupling length constant, *d_i,j_* is the pairwise Euclidean distance between the *i*th and *j*th cells, and *n* is the total number of bipolar cells in the model.

Responses in the model bipolar cell network were then normalized, and output thresholding was then applied by setting values below the threshold equal to zero, and renormalizing the outputs between 0–1. A piecewise nonlinear function (i.e., ReLU) was then applied to the thresholded responses:

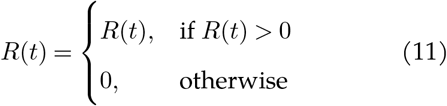

### Mutual information calculations

To compare encoding of past and predictive information, we estimated the amount of information that the neural response at a particular time (*r_t_*) provided about the stimulus at time, *t*′ (**s**_*t*′_), where *t*′ = *t* + Δ*t* using Equation 12. The mutual information was estimated at several different time lags (Δ*t*) relative to the peak of the temporal filter (see (Liu et al., 2021)).

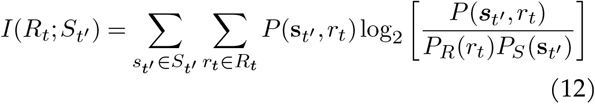

where *P_R_*(**r**) is the distribution of responses in a single cell, *P_S_*(***s***) is the stimulus distribution, and *P* (**s**_*t*′_, *r*_*t*′_) is the joint distribution of stimuli presented at time *t*′ and responses **r** observed at time *t*. In other words, responses were fixed in time, the stimulus was shifted for each time bin, and the mutual information was computed at each of these time shifts (Palmer et al., 2015; Bialek, 2012). These mutual information calculations required converting the spatial dimensions of our stimuli into a single value for each time bin. We did this by first identifying the four spatial regions of the stimulus that were centered over the receptive field. Each of the 16 possible stimulus patterns for those four regions was assigned a value between 0–15.

## SUPPLEMENTARY INFORMATION

**Figure S1.**
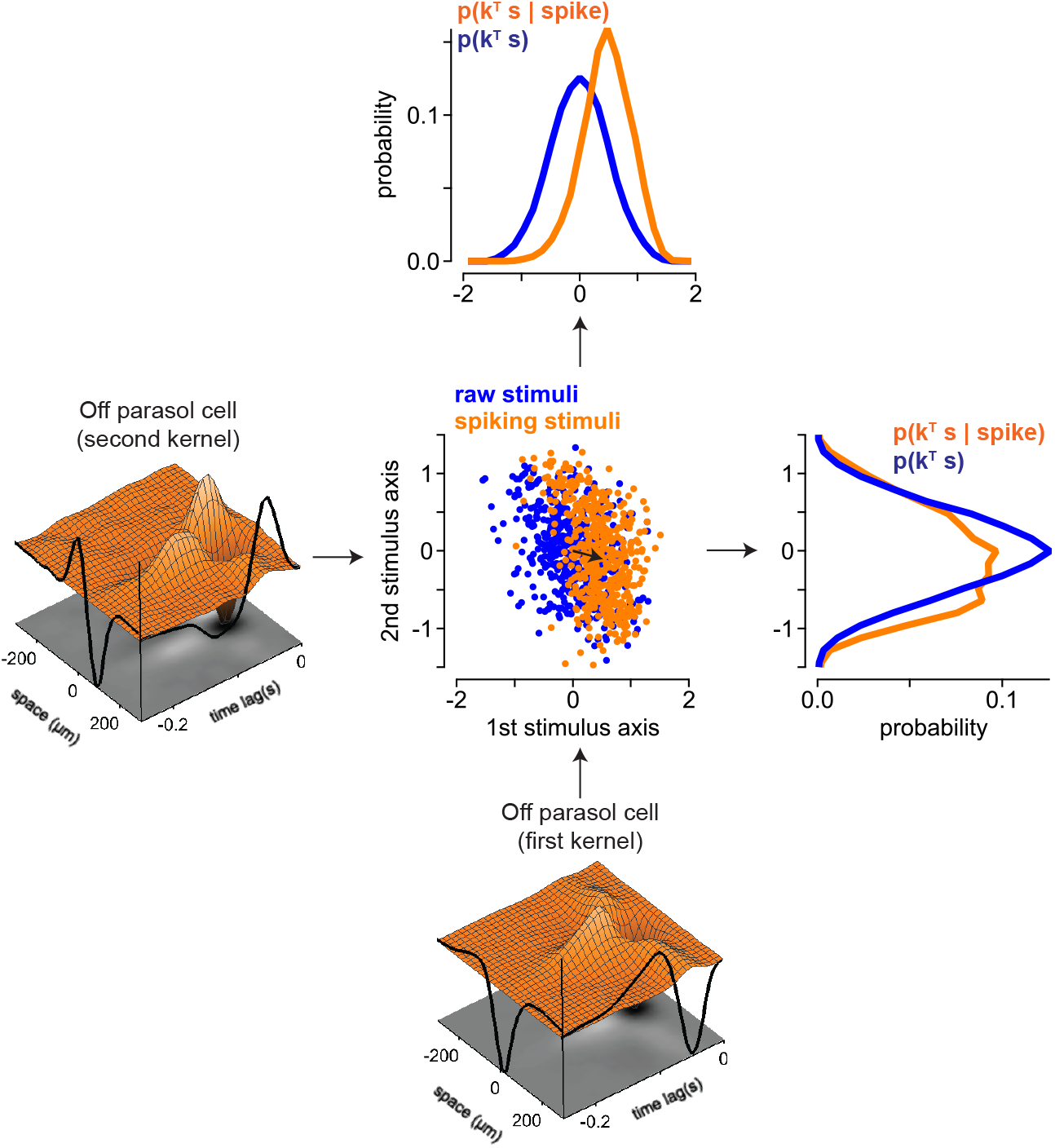
Example of the maximally informative dimensions technique in an Off parasol ganglion cells. *Bottom left*, A two-dimensional stimulus space depicting the raw stimuli (*blue*) and the stimuli that elicited spiking in an Off parasol ganglion cell (*orange*). The black arrow indicates the centroid of the spike-triggered stimuli. The probability distributions for the raw stimuli and the spiking stimuli were computed by projecting the stimuli along the first or second stimulus axis (*top left* and *bottom center*, respectively).

**Figure S2.**
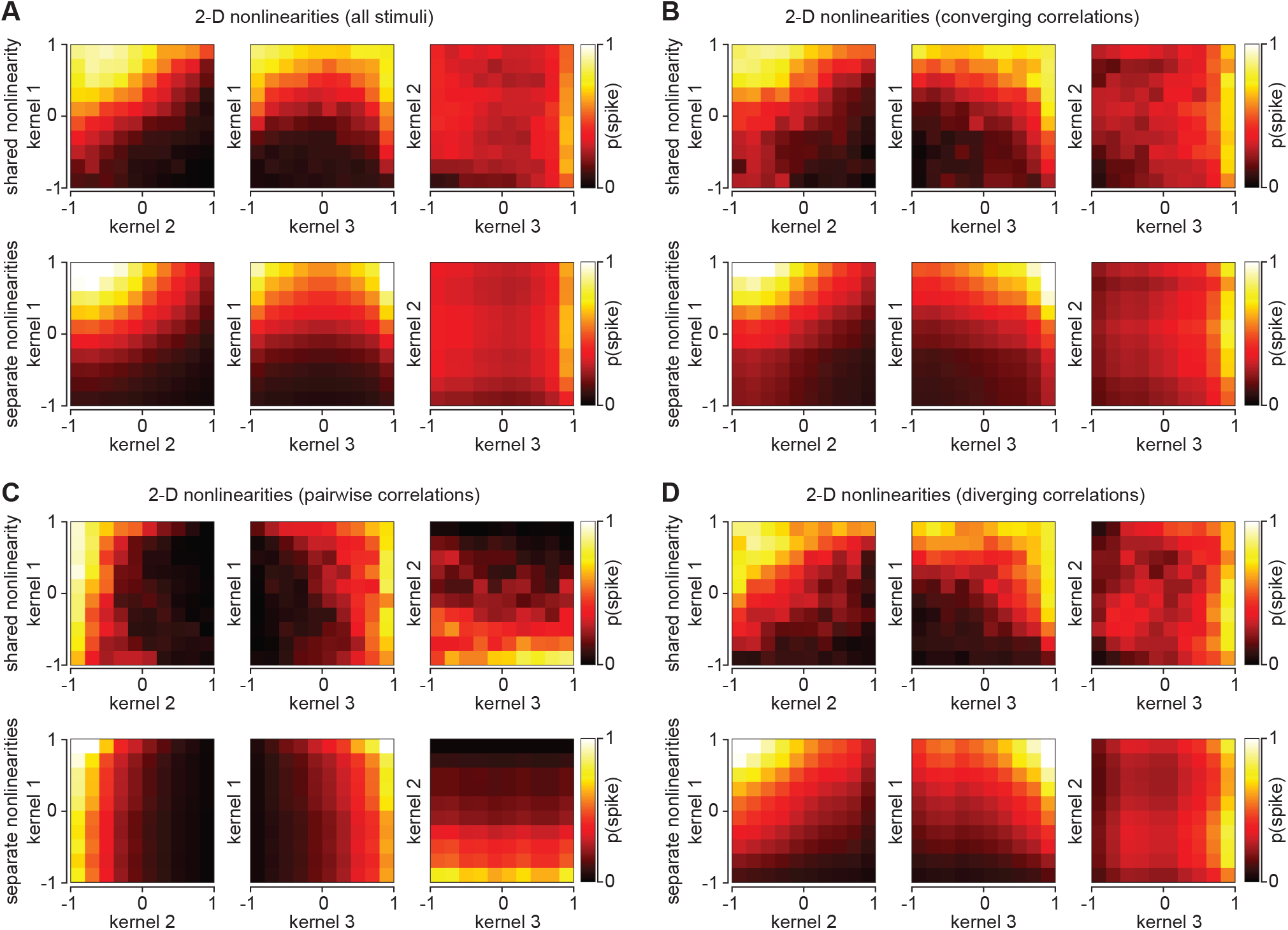
Kernel nonlinearities vary for different classes of spatiotemporal correlation. (A) Two-dimensional non-linearities illustrating the interactions between the individual kernels for an On smooth monostratified cell. The *x* and *y* axes represent the normalized projection of the stimulus onto the individual kernels. The color intensities represent the spiking probability of the cell for a particular location on the interaction map. Two-dimensional nonlinearities are shown for all of the stimulus classes including uncorrelated noise. (B-D) Two-dimensional nonlinearities for converging correlations, pairwise correlations, and diverging correlations in the same cell as (A). The shared and separate nonlinearities differed substantially, indicating that a model in which the kernel outputs passed through separate nonlinearities prior to being combined did not adequately describe the kernel interactions. Further, the shared nonlinearities varied slightly with stimulus class, suggesting that the contribution of the kernels depended on the stimulus correlations.

**Figure S3.**
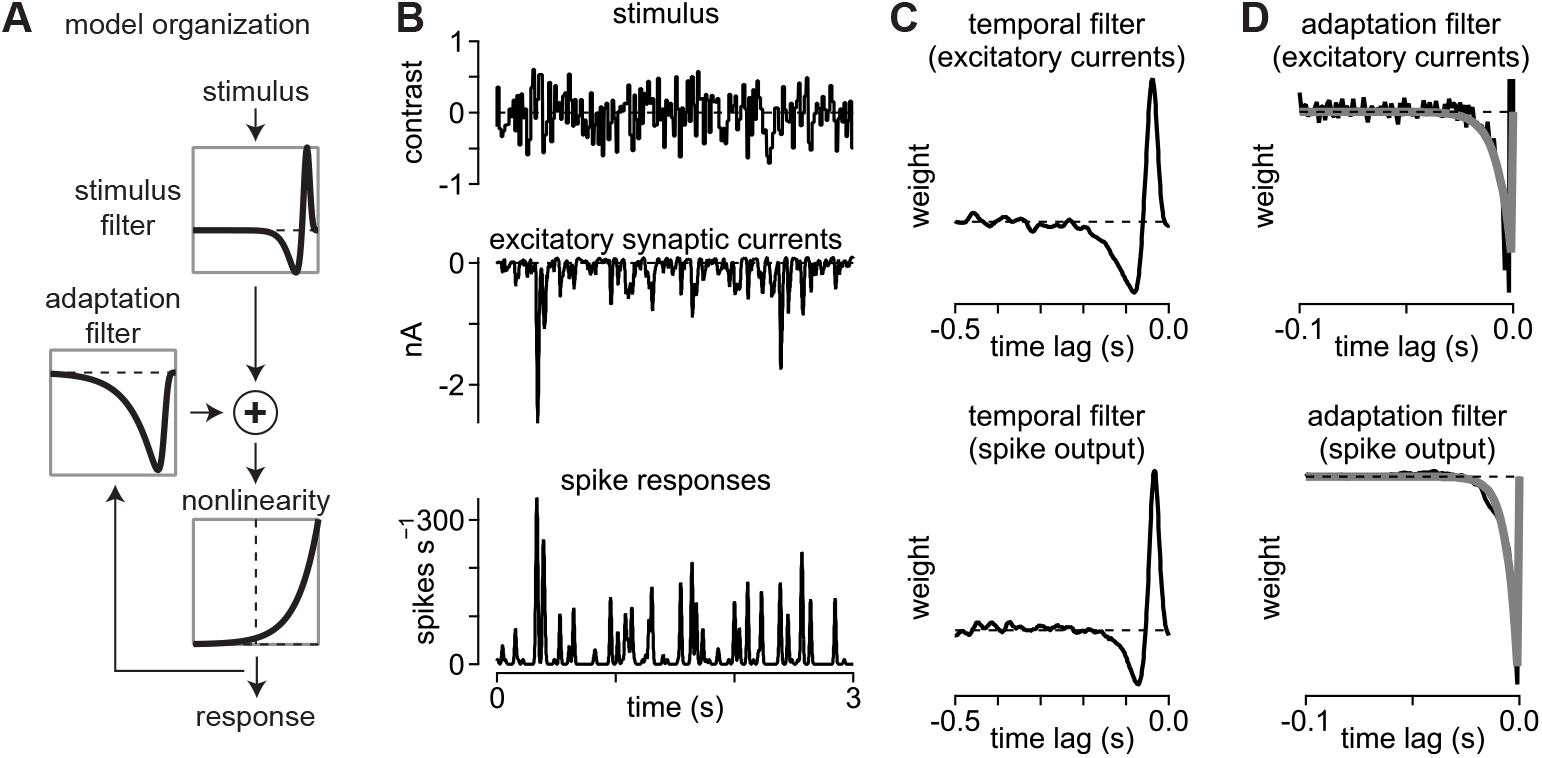
Generalized linear model organization. (A) Gain control in smooth monostratified ganglion cells was estimated using a generalized linear model (GLM). The input stimulus was filtered by a temporal kernel and passed through a nonlinearity. Based on the output history of the model at this stage, an adaptation filter provides feedback to signals prior to the output nonlinearity. (B) Model parameters were estimated by presenting a spatially uniform spot over the receptive field. Spot contrast was drawn randomly from a Gaussian distribution on each time step (*top*). Excitatory synaptic currents were measured to estimate the filtering and adaptation properties of diffuse bipolar cells (*center*) and spike output was also measured in the same On smooth monostratified ganglion cell (*bottom*). (C) Temporal kernels estimated from the excitatory synaptic currents and spike responses in (B). (D) Adaptation filters estimated from the excitatory synaptic currents and spike responses in (B).

**Figure S4.**
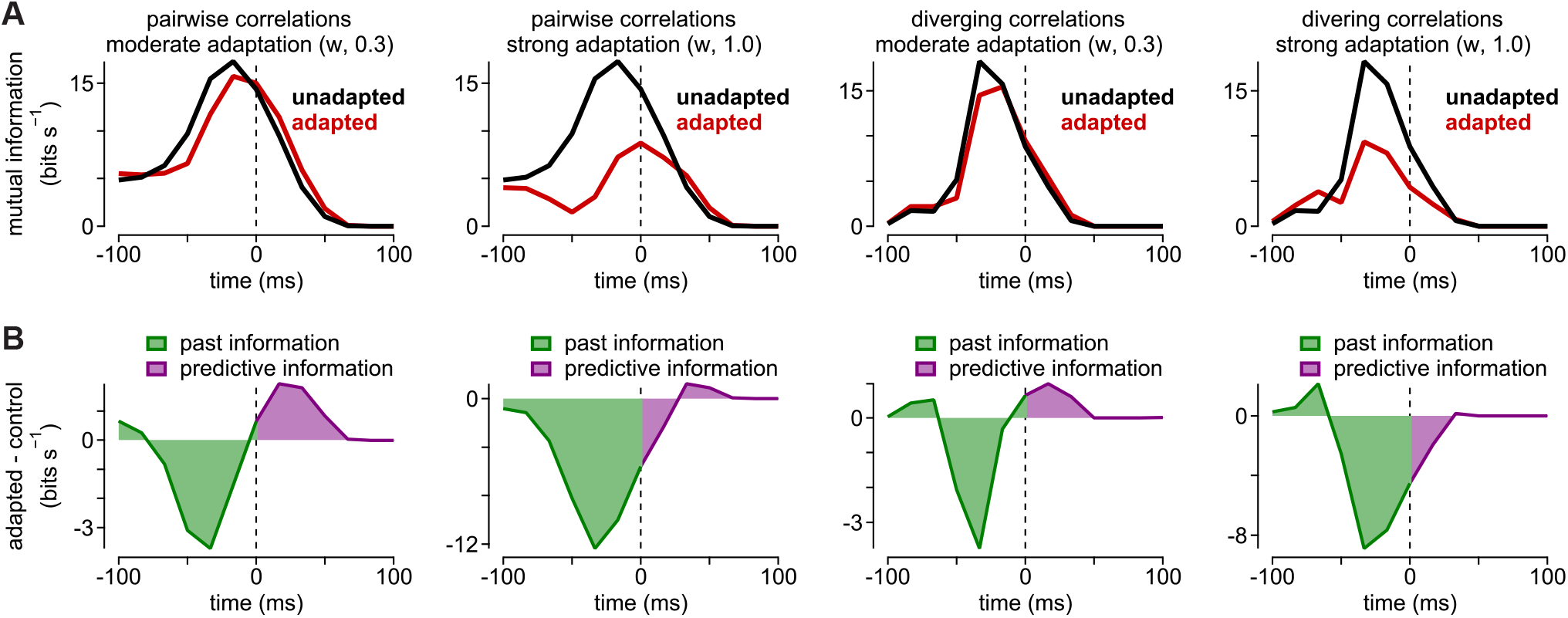
Moderate adaptation improves predictive encoding. (A) Mutual information (*y*-axis) encoded as a function of time lag (*x*-axis) for pairwise and diverging correlations. Curves are shown comparing the model lacking adaptation to models with moderate or high levels of adaptation. (B) Difference curves in which the past (*green*) and predictive information (*purple*) are compared for the adapted versus unadapted curves in (A). Moderate adaptation increased the encoding of predictive information while strong adaptation decreased this encoding.

